# Automatic Sample Segmentation & Detection of Parkinson’s Disease Using Synthetic Staining & Deep Learning

**DOI:** 10.1101/2022.08.30.505459

**Authors:** Bradley Pearce, Peter Coetzee, Duncan Rowland, Scott Linfoot, David T Dexter, Djordje Gveric, Stephen Gentleman

## Abstract

The identification of Parkinson’s Disease (PD) from post-mortem brain slices is time consuming for highly trained neuropathologists, often taking many hours per case. In this study, we demonstrate fully automatic PD detection, from single 1000um regions, from sections spanning from the dorsal motor nucleus of the vagus nerve to the frontal cortex. This is achieved via image processing and statistical methods, with improved accuracy demonstrated when using machine learning. Digitised stained brain sections were processed via a deep neural network to produce re-coloured, or ‘synthetically stained’, images which were then filtered and passed to a secondary network for classification. We demonstrate state-of-the-art PD detection (>90% accuracy on single 1000um regions), with the ability to perform binary classification on high resolution sections within minutes, in addition to demarcating regions of interest to the pathologist for manual visual verification.

**Executive Summary:** The identification of Parkinson’s Disease (PD) from post-mortem brain slices is time consuming for highly trained neuropathologists, often taking many hours per case. Accurate classification and stratification of PD is critical for the confirmation that the brain donor suffered from PD and to maximise the potential usefulness of the brain in research studies to better understand the causes of PD and foster drug development.

Parkinson’s UK Brain Bank, at Imperial College London, has produced a dataset containing digitised images of brain sections immunostained for the protein alpha-synuclein (*α*-syn), the pathological marker of PD; along with control cases from healthy donors. This dataset is much larger (over 400 cases), more consistent, and of higher quality (all have been stained with the same protocol and imaged within the same laboratory) than has been documented elsewhere in the literature; including those found in a meta-analysis study on detection of neurological disorders containing over 200 papers (Lima et al., 2022).

The project team, consisting of neuroscientists and subject matter experts from: Imperial, NHS AI Lab Skunkworks, Parkinson’s UK, and Polygeist have undertaken a 12 week project to examine the possibility of producing a Proof-of-Concept (PoC) tool to automatically load, enhance and ultimately classify those brain sections containing *α*-syn. The initial focus of the project was to make a tool that could make a biomarker of PD, *α*-syn, more visible to the pathologist; saving time in searching for the protein manually. This goal was quickly reached, producing a tool that could ‘synthetically stain’ the *α*-syn, marking regions of interest in a high-contrast bright green, making them quickly identifiable for the pathologist. Statistical analysis of the synthetically stained images showed that very few regions in the control group were stained compared to the PD group, raising the possibility that an automatic classifier could be developed, which became a stretch goal for the project.

A bespoke neural network model was designed that processed the synthetically stained segments of each immunostained section and produced a binary judgement (whether a segment contains PD pathology or not). The model achieved >90% sensitivity for PD detection, much higher than is reported for neuropathologists (~60% sensitivity when searching for *α*-syn patches across all stages, Signaevsky et al., 2022). While expert raters are still more precise (~6% better than the model), the model performed ~20% better than expert raters when considering precision and recall.

The key output of the project is an open-source PoC tool that can automatically classify PD from digitised images of brain sections with accuracy that is approaching viability for real world applications. An MIT Licensed code repository has been released, containing all of the model development code, along with associated documentation, to allow others to build on the project team’s work. This report summarises the scientific and engineering process undertaken through the development of the PoC tool.

## 0 Extended Summary of Sections

The **Technical Introduction** section gives a brief background on the protocol of *α*-syn synthetic staining process and a brief description of prior machine learning and image processing efforts on automatic identification. The latest studies that were found required expert raters to identify regions of interest in biopsies of slides before they were passed to machine learning algorithms. This is likely due to having a low sample size and comparably many more patches containing no pathology than those that do. This section describes our two predictions. First, automatically synthetically staining regions of interest will allow for visual identification of regions of interest (and hence automatic dataset generation). Second, our machine learning architecture will achieve a high accuracy binary identification of PD when classifying this enhanced dataset.

The **General Methods** section gives a detailed description of the process followed to replicate our findings, including the materials used, the workflow, and the machine learning architecture. It also documents the configuration parameters used in the processing pipeline along with the process of generating fake imagery for testing purposes. It documents the processing pipeline: taking an initial file, breaking it into patches so it can be loaded onto the GPU, staining those patches using an autoencoder, filtering those patches to identify regions which were stained, and then training the machine learning model to classify those filtered stains as either PD or control.

The **Machine Learning for Automatic PD Classification** section details the machine learning architecture used. This includes a base network, EfficientNet, which has previously been shown to classify biological imagery with high accuracy (Munadi et al., 2020), and then additional layers specific to classifying synthetically stained imagery.

The **Results** section gives a detailed description of outputs of the synthetic staining, with example comparisons of those outputs to original brain sections, showing how PD-like pathology is more easily identified. It also shows the high frequency of patches in the PD group that were identified as containing staining, along with example patches showing the visible differences between the groups. The training and validation accuracy graphs are presented, along with the performance of the final system, using a variety of metrics. The results validate the assumptions made in the creation of the processing pipeline. This pipeline involves creating a known signal (synthetically stained proteins), filtering it for the initial validation that the signal differentiates the groups (detecting that signal with a weak filter, producing stained regions for both the PD and control groups), and then building a classifier that can automatically build the relevant filters to perform binary classification (binary classification of PD or control group). From these results, we conclude that the processing pipeline is sensitive to *α*-syn, can effectively discriminate the PD and control groups on the basis of this signal, and can be produced programmatically from source images, configuration files and fitting using back propagation.

The **Discussion** section gives a general overview of the results. This includes the viability of the synthetic staining technique, along with an explanation of why certain artefacts remain, and how the research may be developed further in the future. The discussion describes how these technologies may be used to produce finer grain assessment than binary classification, such as providing a severity score for PD, or providing automatic Braak staging.

The report closes with a **Next Steps** section which recommends further work to maximise the impact and applicability of these results for histopathologists engaged in research on brain tissues.

The **Conclusion** section summarises the work described through this report and the outcomes which it enables. A **Glossary of Terms**, containing a list of definitions and abbreviations follows at the end of the References section.

## 1 Technical Introduction

Accurate diagnosis of Parkinson’s Disease (PD) is essential to furthering our understanding of the pathological causes of PD which will ultimately lead to the development of more effective therapies. Post-mortem neuropathological diagnosis of PD is labour intensive, requiring manual microscopic examination of stained brain sections and adherence to a number of diagnostic protocols and staging systems. The predominant staging system, Braak staging (Braak et al., 2003) defines six stages, progressing from Stage I (the beginning of disease, in the dorsal motor nucleus of the vagus nerve), to Stage VI (the apex of the disease, within the neocortex). These protocols are the subject of active research, with variable inter-rater reliability between expert raters, when staging pathology (0.4 < Krippendorf’s *α*^1^ < 0.6, for review and comparative analysis, see Attems et al. 2021). These protocols share in common the identification of *α*-syn pathology at various sites within the nervous system including olfactory, neocortical and subcortical structures, and the brainstem (Attems et al., 2021; Alafuzoff et al., 2009; Braak et al., 2003; McKeith et al., 1996), along with reports on clinical presentation. Moreover, high-fidelity quantitative stratification of the degree of PD pathology is not yet available, due to the manual nature of staging,

Lewy Bodies (LBs) are largely constituted by *α*-syn protein, which can be stained via Immunohistochemistry (IHC), with commercially available antibodies with Diaminobenzidine (DAB) detection (for comparative study, see Croisier, et al., 2006). Visual identification of both intracytoplasmic and extracellular *α*-syn can be made using light microscopy (using Aperio scanners) at 20x objective (~200x visual) magnification (see Crosier, et al., 2006; Beach, et al., 2018). Despite the viability of obtaining digitised imagery of samples containing stained *α*-syn the authors are not aware of any high-sensitivity, automated PD classification and stratification algorithms utilising machine-vision and machine learning.

There has been extensive research into automatic diagnosis of PD in patients using machine learning, including assessment of patient speech and gait (for comprehensive review, see Mei, Desrosiers & Frasnelli, 2021); however, these studies usually rely on post-mortem neuropathological diagnosis to assess accuracy with no ability to stratify stages of pathology. There has been some success automatically classifying antemortem biopsies using machine vision and convolutional neural networks; Signaevsky et al. (2022) report ~69% accuracy in a forced-choice binary classification of IHC stained PD biopsies using a Convolution Neural Network (CNN) algorithm; however, the biopsies still required manual *α*-syn Region-Of-Interest (ROI) labelling, with these labels exhibiting poor inter-rater reliability (error ±13%). The true sensitivity of these systems are also poorly understood: in the latter study, samples were taken from 42 PD patients and 14 controls. Many more samples would be required to assess the sensitivity of the CNN algorithm relative to its hypothetical performance when compared to object classification tasks with many samples (see Redmon, Divvala, Girshick & Farhadi (2016) for benchmarking methodology).

For the development of high accuracy, automatic classification of PD, a number of material advancements are required. Firstly, a dataset is required that has homogenous variance for both the PD and control samples (in sampling and staining protocol, and imaging etc.), including a large enough sample size to yield statistical power for algorithm design, and classification performance analysis. Secondly, a methodology to automatically and consistently identify ROIs for classification, to both the pathologist/expert rater (to verify those ROIs), and to provide a uniform classification criterion to a secondary machine learning model (high inter-rater reliability).

In this study the Parkinson’s UK Brain Bank, at Imperial College London, provided samples of brain tissue (hereafter referred to as slides) from 401 cases (301 PD cases). All slides were acquired and digitised using the same methodology, and are described in more detail herein. The study aimed to produce an algorithm that would reduce the time-to-identification for expert raters to identify *α*-syn stained bodies, which leads directly to a neuropathological diagnosis of either PD or associated Lewy Body Diseases (LBDs). To do this, we aimed first to identify the *α*-syn bodies, which vary in size, chromaticity and texture, and then differentiate those bodies against the rest of the slide contents, by increasing their visual saliency through recolouring *or “synthetic staining”* (no saliency metrics are discussed or employed here, see Itti & Koch (2001)). This involves building a model that is aware of the texture, shape and colour variations of the stained *α*-syn, yet insensitive to bodies of similar shape and colour (such as grains of neuromelanin) that are present in neurotypical case samples, and would be processed equally using image processing such as simple masking (see Law et al. 2017).

Zhang et al (2017) proposed an autoencoder based neural network (referred to here as iDeepColour), which accepts grayscale images along with a chromaticity mask as input, and produces RGB output (with reduced spatial resolution). iDeepColour is trained to be a general purpose, texture and spatially aware ‘flood fill’ algorithm (for discussion on texture aware floodfill, see He, Hu & Zeng, 2019). As *α*-syn bodies vary in size, chromaticity and texture, we predicted that a conservative chromatic mask would only partially identify *α*-syn bodies (and other, erroneous, bodies of similar chromaticity). However, we predicted that the output of iDeepColour given the chromatic mask (with a grayscale version of the slide), would be differentially filled regions, with proportionally more *α*-syn stained than erroneous bodies (as can be seen for natural imagery in Figure 1.1).

**Figure 1.1.**
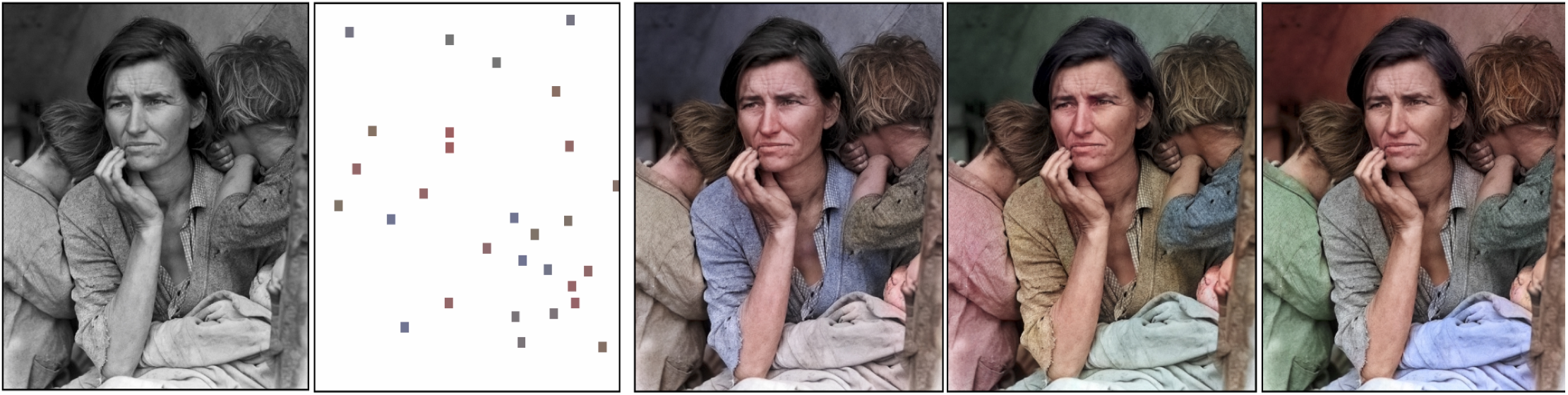
Masking and recolouring of natural images using iDeepColour *(from Zhang, et al*., *2017)* **Left**. Grayscale scene, provided as input along with a chromatic mask (white denotes that the entire image is to be coloured, coloured squares are ‘hints’ to the neural network (iDeepColour) performing the colouration (flood fill); **Right**. Output of different runs of iDeepColour with varying mask chromaticities.

To quantify the effectiveness of *α*-syn isolation through synthetic staining, we proposed segmenting the slides into patches, and retaining those that contained stained bodies, as in Signaevsky et al. (2022). This allowed for a sign test (testing whether more patches were retained proportionally from the PD group than Controls) as well as further analysis using neural networks designed for image processing.

Signaevsky et al. (2022) utilised the Inceptionv4 architecture (with 42M trainable parameters, see Szegedy et al., 2017), with a final fully connected layer for binary discrimination. Inceptionv4 has demonstrated little benefit in binary classification of medical imagery previously; Yang et al., (2021) demonstrated that EfficientNet performed comparably with less than 100,000 parameters (using the b0 variant of the network) when classifying COVID-19 patient x-ray scans. Munadi, Muchtar, Maulina & Pradhan (2020) also demonstrate high accuracy (~95%) using a pre-trained EfficientNet model for image enhancement and tuberculosis detection. In these studies large volumes of data can be processed quicker when utilising models with fewer parameters, increasing the number of training epochs available within a fixed time.

We predicted that EfficientNet would be a viable candidate for binary classification training of PD pathology from synthetically stained *α*-syn patches, given the two-week sprint cycle that was available for model development, training and validation. We predicted that given the absence of *α*-syn bodies within the control group, erroneously stained patches within the control group would have a different qualia with respect to shape, texture and size; these differences are good candidates for filter development. We expected that a trained EfficientNet model would have significantly higher classification rates of ‘PD’ for the PD group than the control group, with high specificity and recall.

Here we present the development, preprocessing, training and validation procedure for automatically detecting PD via post-hoc classification of synthetically stained *α*-syn patches. Full software source code, including engineering assurance, documentation and ancillary project outputs can be found on the NHS Transformation Directorate public github repository at: https://github.com/nhsx/skunkworks-parkinsons-detection.

## 2 General Methods

### Materials

These experiments utilised digitised images provided by the Parkinson’s UK Brain Bank. Images at 20x magnification acquired using Aperio scanners were utilised. Images were in Aperio (.svs) format, with each pixel width and height spanning .504µm; typical tissue area was ~6.6cm² per sample. Samples were sections of human tissue, ~7µm in thickness, taken post-mortem from between 8 and 11 regions within the Central Nervous System (CNS) per case. Brains were fixed in formalin and sections of brain tissue were subsequently stained with an IHC protocol targeting *α*-synuclein protein with Diaminobenzidine (DAB) intensification and a subsequent Hematoxylin counterstain. Relevant slide site descriptions can be seen in Table 2.1.

**Table 2.1.** Slide site index, associated anatomical region, and Braak stage when pathology is present.

| Slide Site Index | Anatomical Region | Associated Braak Stage |
| --- | --- | --- |
| 1 | Frontal Cortex | 6 |
| 2 | Cingulate Cortex | 5 |
| 4 | Temporal Cortex | 4 |
| 15 | Substantia Nigra | 3 |
| 16 | Locus Coeruleus | 2 |
| 17 | Dorsal Motor Nucleus of the Vagus | 1 |

Samples were from 401 cases, 301 which had been classified as containing pathology associated with PD, images totaling 4154 were assessed (3185 PD samples, 969 from controls, between 8-11 per case).

All experiments were performed using executable computer code running atop generally available workstation hardware (System); using an AMD Ryzen Threadripper 2950X (32-core @ 3.75GHz), NVIDIA GeForce RTX 3090 (24GB VRAM), and 64GB of core memory. The system utilised Linux 5.18.7, CUDA 11.7 (r11), PyTorch 1.12.0, and the NVIDIA APEX extensions for PyTorch (a0f5f3a). All other software configurations, and installation instructions can be found on the public GitHub repository at https://github.com/nhsx/skunkworks-parkinsons-detection (herein, Repository). Some of the software dependencies that were utilised were subject to third party licence agreements (graphics and compute drivers for the RTX 3090), which were accepted and utilised.

A pretrained Caffe model file was utilised from Zhang et. al. (2017) for image colourisation, along with executable code for the iDeepColour convolutional neural network which can be found within the Repository.

### Preprocessing

All slides were in 24bit Aperio sRGB format with equal exposure and transformed from RAW sensor space by the Aperio scanner (see Olson, 2013). Images were loaded into memory using a configuration protocol (available within the Repository) and the SlideIO library (Melnikov & Popov, 2021). In-memory images were interpolated using bilinear interpolation and were compared to QuPath (Bankhead, et. al., 2017) with minimal differences in interpolated value (<1% RGB). All images were loaded with 4 pixel downsampling, each pixel spanning approximately 4µm² in area. Each 8-bit channel (0-255 intensity) was converted (scaled without in-frame normalisation) to a 16-bit floating point value between 0-1, for use within Torch with CUDA (see the Repository for details).

### Synthetic Staining Procedure

Each slide was loaded into memory and segmented into 512px² patches (spanning ~1000µm), see Figure 2.1. A binary mask was produced for each patch, produced by thresholding the blue colour channel, which denoted RGB pixels with a blue component less than 0.196 (Mask M is True where *b*<0.196 otherwise it is False). Within our dataset this threshold was found to identify *α*-syn protein, as well as neuromelanin and cell nuclei, and other darker pigmented bodies sensitive to the staining wash. This mask was passed along with the G-channel of the patch (grayscale image) to the iDeepColour neural network, which included a resizing step to 256px². This network processed each patch to produce a colourised (synthetically stained) patch where mask areas were filled with the deep-priors, texture-aware filling algorithm as described in Zhang et al (2017). Small areas of erroneously masked, high-frequency texture are ignored by the algorithm with more uniform areas associated with *α*-syn protein filled. The resulting output patches were reassembled into a coherent image and inverted to enhance the visibility of the colourisation (I_out_ = 1 − I_in_, where I_in_ is the input image). These stained images were saved to disk, as lossless 32-bit PNG images. A schematic of the procedure can be found below, in Figure 2.2.

**Figure 2.1.**
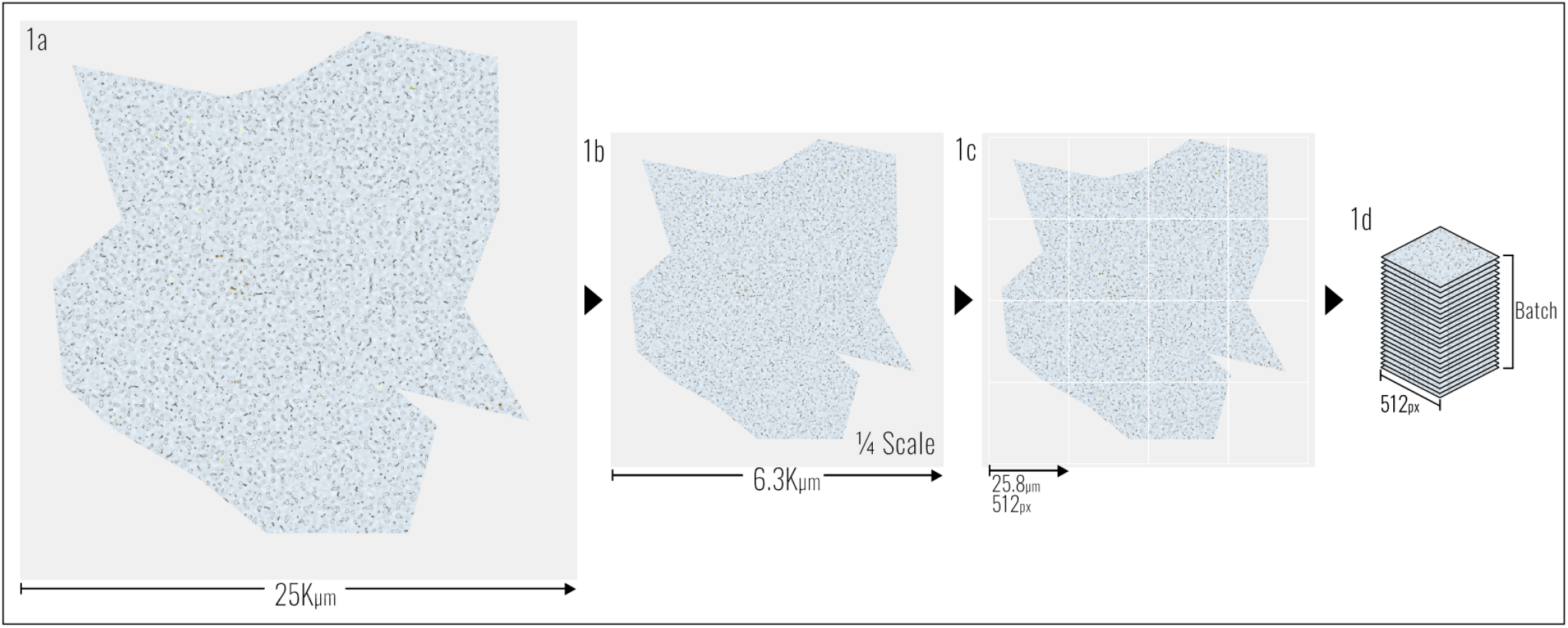
Loading and segmentation process from .svs to Torch tensor. Synthetic image data presented. *(illustration only, not to scale)* **1a**. Synthetic slide, in .svs format at native resolution spans approximately 55K pixels, at ½ micron per pixel; **1b**. When loaded into memory as an RGB array, the image is downscaled to ¼ resolution; **1c**. Patches are specified as 512 pixels wide and segmented; **1d**. Patches stacked as a batch for input into Torch handled neural networks.

**Figure 2.2.**
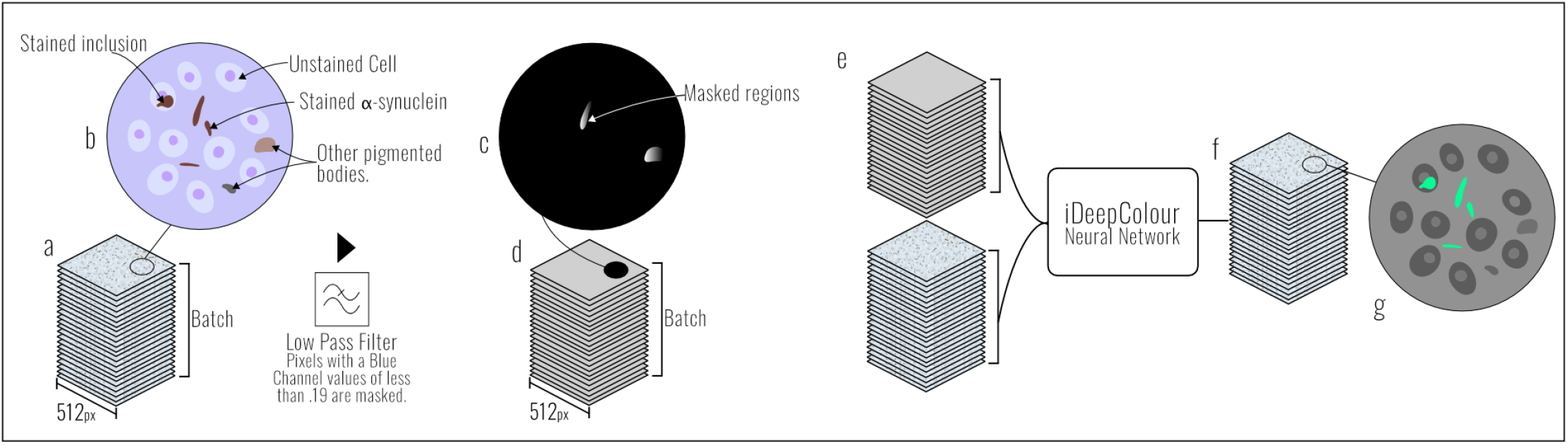
Masking and synthetic staining of slide patches using iDeepColour. *(illustration only)* **a**. Patch stack produced during data loading; **b**. Schematic of cell and staining types, showing confusable bodies and targets for synthetic staining; **c**. Low-pass filter, using blue threshold *b* <.196 produces binary mask with some targets and confusables masked; **d**. The binary mask batch is produced by filtering the initial tensor stack; **e**. The binary mask batch and grayscale batch of slides is passed to iDeepColour for deep-prior aware flood-filling; **f**. Output batch of synthetically stained patches; **g**. Target regions are coloured bright green, with confusion regions and cells remaining grey.

### Runtime Statistics

While the executable code for these experiments was neither designed with runtime performance in mind, nor profiled or optimised, wall-time metrics of the processing pipeline were recorded, and directly relate to the hardware specified above.

### Machine Learning for Automatic PD Classification

Segmentation of Synthetically Stained Images for Machine Learning

Synthetically stained images for each case’s dorsal motor nucleus of the Vagus (slide index 17) were re-traversed in 512px² patches, as in Figure 2.2. The patches were tested for the degree of synthetic staining by comparing the red and green channels. If at least .5% (approximately 3 pixels) had green staining (20 intensity values greater than the red channel, a measure of luminance in this case), then the patch was written to disk as a 512px², 32-bit PNG image. Patches were separated into their associated PD and Control groups, and further separated into training and validation datasets for machine learning training and testing. The training set comprised 75% of all slides.

### Machine Learning Architecture

The model utilised a variation of the EfficientNet-B7 (EffNet) model proposed by Tan & Le (2019). A number of additional layers were added to allow for robust binary classification. These included: a concatenation layer consisting of fully connected nodes (torch.nn.Linear), dropout layers (30% of values dropped per layer) to encourage robust filter generation, and a sigmoid layer to produce confidence scores between 0-1 with the mean just-noticeable-difference (JND) at confidence=.5. A preprocessing layer was added to transform inputs from a core memory array (512px² x RGB in size) to EffNet’s input layer shape (Torch HalfTensor, 299px² RGB). A network schematic can be seen below in Figure 2.3; a full implementation can be found in the Repository. This network will be referred to here as ‘PDNet’.

**Figure 2.3.**
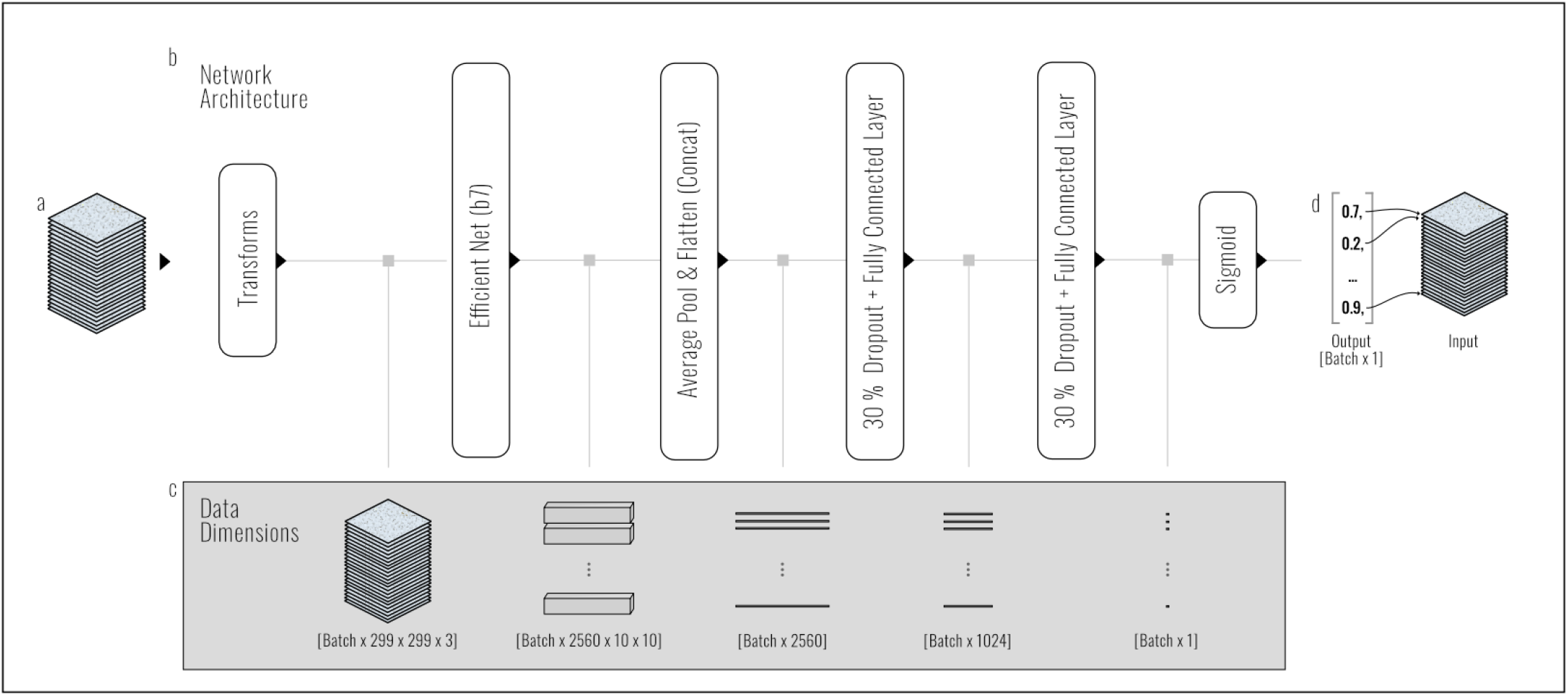
PDNet Architecture & Data Flow. **a**. Patch stack produced during data loading; **b**. Network architecture of PDNet, with data flow left to right; **c**. Schematic of data dimensions: data starts post transformation to EffNet’s input size (299px² x RGB), data dimensionality is reduced by each layer until a single value is output; **d**. Output, with example mapping to input stack.

### Training Configuration and Procedure

Training inputs underwent an additional augmentation step, using Torch’s transformation library; these included a random horizontal flip, and random image rotation both in 50% of patches per epoch (for details, see Pointer, 2019); these transformations were not applied to validation (test data). PDNet was trained for 500 epochs, using stochastic gradient descent with a learning rate of *lr*=0.0125 and momentum of *m*=0.9. Binary cross entropy, with logit loss was utilised as a loss criterion (for details, see Bishop, 2006). Patches were split into 32 image mini-batches. Training and test patches were passed to PDNet with the output resulting in a floating point value between 0 and 1 for each input patch. That output, taken to be a confidence score, was converted to a boolean classification by assuming any value other than 0 was a PD group classification (L= *f* (C > 0, G), where L is our loss, *f* is our loss function, C is our confidence score for the patch, and G is our group, with 1 denoting the PD group and 0 the control). This configuration was designed to minimise overfitting, by punishing any non-0 output in the control group. The loss was then scaled and backpropagated to the network (see Hayin, 2004), using the optimiser functions provided by APEX (see Repository for details), for each 32 patch batch.

### Testing & Validation Procedure

During the validation phase the same loss metric and binary classification accuracy was computed for the validation dataset (not seen by the network for model fitting). An additional phase of testing was performed where confidence scores were produced for all PD and control patches in the validation dataset; these were then thresholded using a sliding threshold between 0 and 1 to calculate the sensitivity, specificity, precision and accuracy statistics, and weighted F measures. These statistics were calculated on a per patch level, rather than per case. As above, the training set comprised 75% of all slides, and 25% were reserved for validation.

### ROI Annotation

Each synthetically stained slide of index 17 (dorsal motor nucleus of the Vagus) was loaded and segmented (as described above), then passed to PDNet for classification via the validation procedure above. If PDNet produced a confidence score of C > .95 for that patch then that region in the stained image was annotated by setting the border pixels of that region to red ([r = 1., g = 0., b = 0., a = 1.]). These annotated images were then written to disk as PNG files.

### Synthetic Data Generation

To test the software pipeline we generated synthetic image data designed to mimic the shape and colourimetric qualities of the slide stimuli. These synthetic images were not designed to be visually comparable to the ecological imagery, but to mimic the shape and form, with differentiation, to allow for testing. These images were procedurally generated using the Perlin worms algorithm (for examples of how this technique has been applied to other medical image generation see Dustler et al., 2015). Three Perlin planes were generated for each synthetic image, using 10 octaves and a random seed; the seeds were conjoined, allowing a single seed to be used for each image. The Perlin planes were sliced with band pass filters (see the Repository for implementation details), to produce a cellular-like mask, which was flood-filled with various RGB values sampled from the slide dataset. Finally, a randomised convex polygonal mask was used to produce a slide outline with the outer regions being set to maximum white (RGBA=1) as in the slide samples. For schematic and examples see Figure 2.4 below.

**Figure 2.4.**
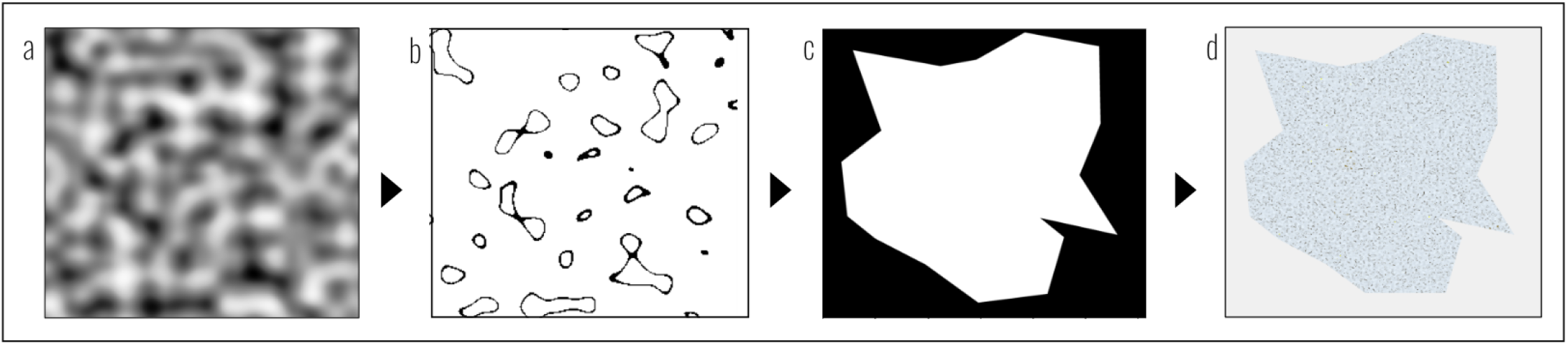
Synthetic Slide Creation Workflow. **a**. Perlin noise planes are generated; **b**. Planes are bandpass filtered, with mask regions assigned RGB values and composited together; **c**. Random convex hulls are generated and used as a binary mask to create tissue-like boundaries; **d**. Output, RGB image with cellular bodies of varying RGBs within the tissue-like boundary.

## 3 Results & Analysis

### Synthetic Staining & Post Processing

Figure 3.1 below also shows *α*-syn bodies identified in a texture-aware fashion, that are not fully filled by a simple mask, and how some highly textured bodies which were erroneously masked are unfilled. This is evidenced by the observation that the binary mask is jagged (see b in Figure 3.1), but the stained output is smooth and filled. Each slide from the 100 control cases and 301 PD cases were successfully synthetically stained. Figure 3.2 for a control and PD case demonstrates the typical differences between the groups. Synthetically stained imagery appears dark (due to inversion) with stained proteins appearing bright green. Visual assessment *prima facie* shows significantly more synthetic staining for the PD group; *α*-syn is synthetically stained green, see Figure 3.3 for magnified comparisons.

**Figure 3.1.**
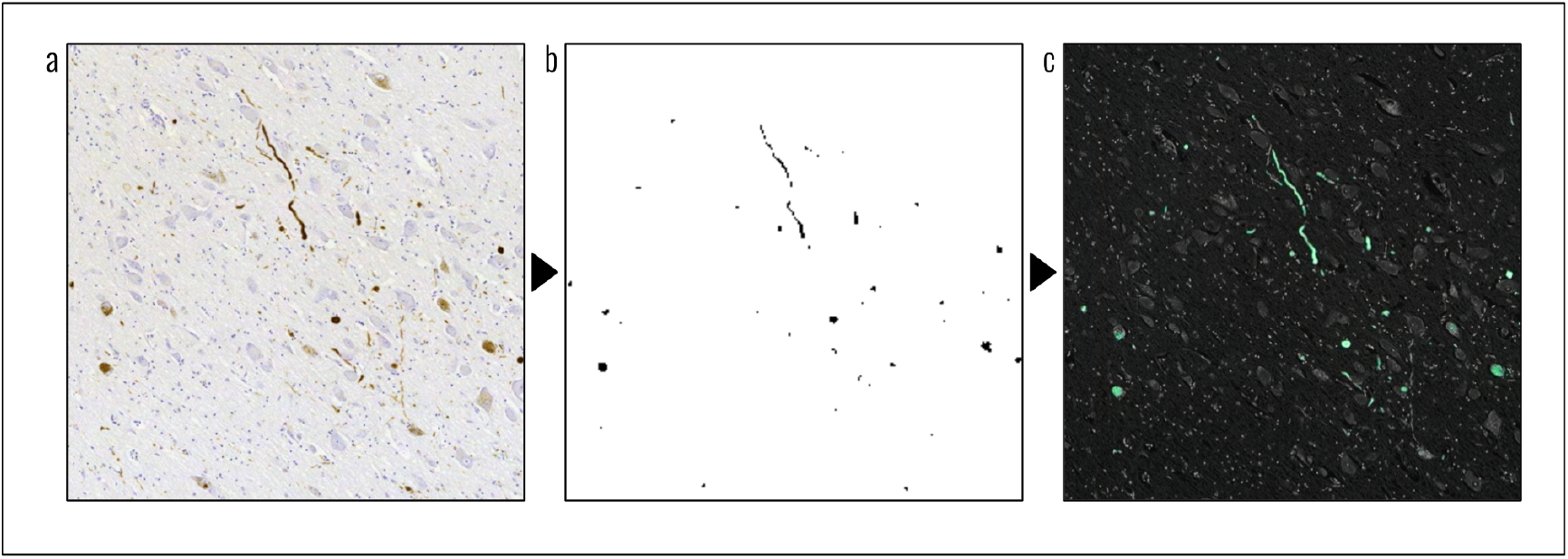
Slide masking and staining process, demonstrating the texture aware flood-filling provided by iDeepColour. **a**. Original slide region, containing target protein. **b**. A conservative mask, showing some specificity for the protein, with erroneous bodies. **c**. Smoothly filled protein, with some additionally unmasked staining, and dodging of highly textured masked regions.

**Figure 3.2.**
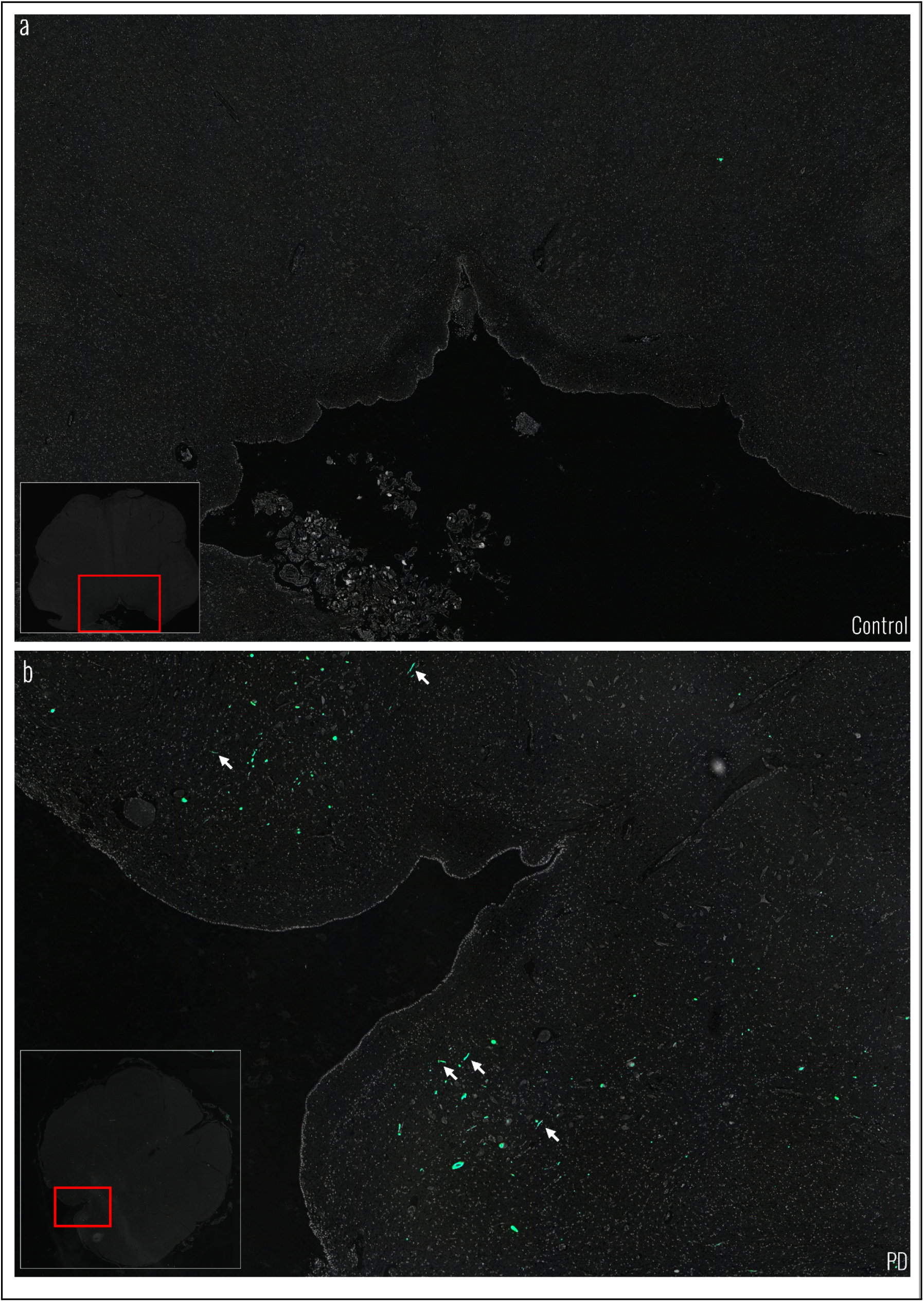
Slides of synthetic staining of the dorsal motor nucleus of the Vagus for a PD and control case, showing LBs and erroneous stains. *(Gamma Corrected)* Synthetically stained slide region, stained bodies in green; **a**. Control case; **b**. PD case; see red cutouts for ROI magnified. Arrows indicate example LBs.

**Figure 3.3.**
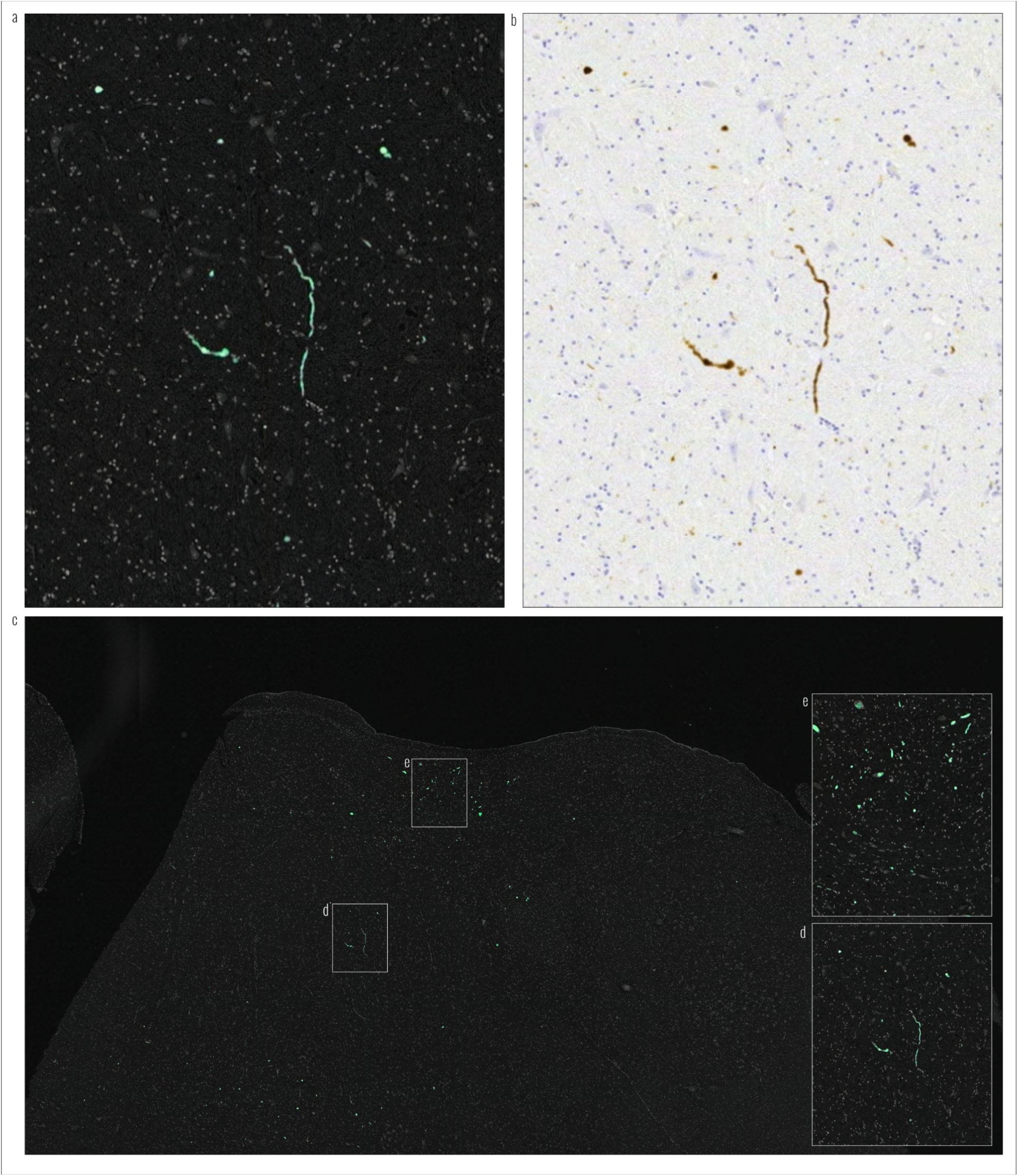
Magnified raster of the DMNoV slide of a PD case. *(Gamma Unchanged)* **a**. Digitally magnified synthetically stained slide region, showing *α*-syn proteins stained green; **b**. Original slide region showing *α*-syn proteins; **c**. Synthetically stained slide digitally downscaled, demonstrating ease of staining identification. **d**,**e**. Denotes regions that are inset and magnified in the right of the pane.

Slides of the dorsal motor nucleus of the Vagus (slide index 17) were processed for each group to test proportionally differential synthetic staining, see *‘Segmentation of Synthetically Stained Images for Machine Learning’* described above (all PD cases were expected to contain some staining in this region, and all controls were expected to be absent of staining). Of the 216,008 patches processed (157,943 PD), 2,582 patches passed our threshold filter compared to 762 control patches; with a median of 3 patches per PD case and 0 patches per control case, see Figure 3.4 for box plots that show the frequency of patches that passed filtering for each group, showing a greater spread and density in the PD group. As the distributions were not normally distributed (see tails in Figure 3.4), we performed a non-parametric Kruskall-Wallis test to determine the probability of committing a Type I error^2^; which showed significant differences between the number of patches filtered between the groups H(1,461) = 25.476, p<.001.

**Figure 3.4.**
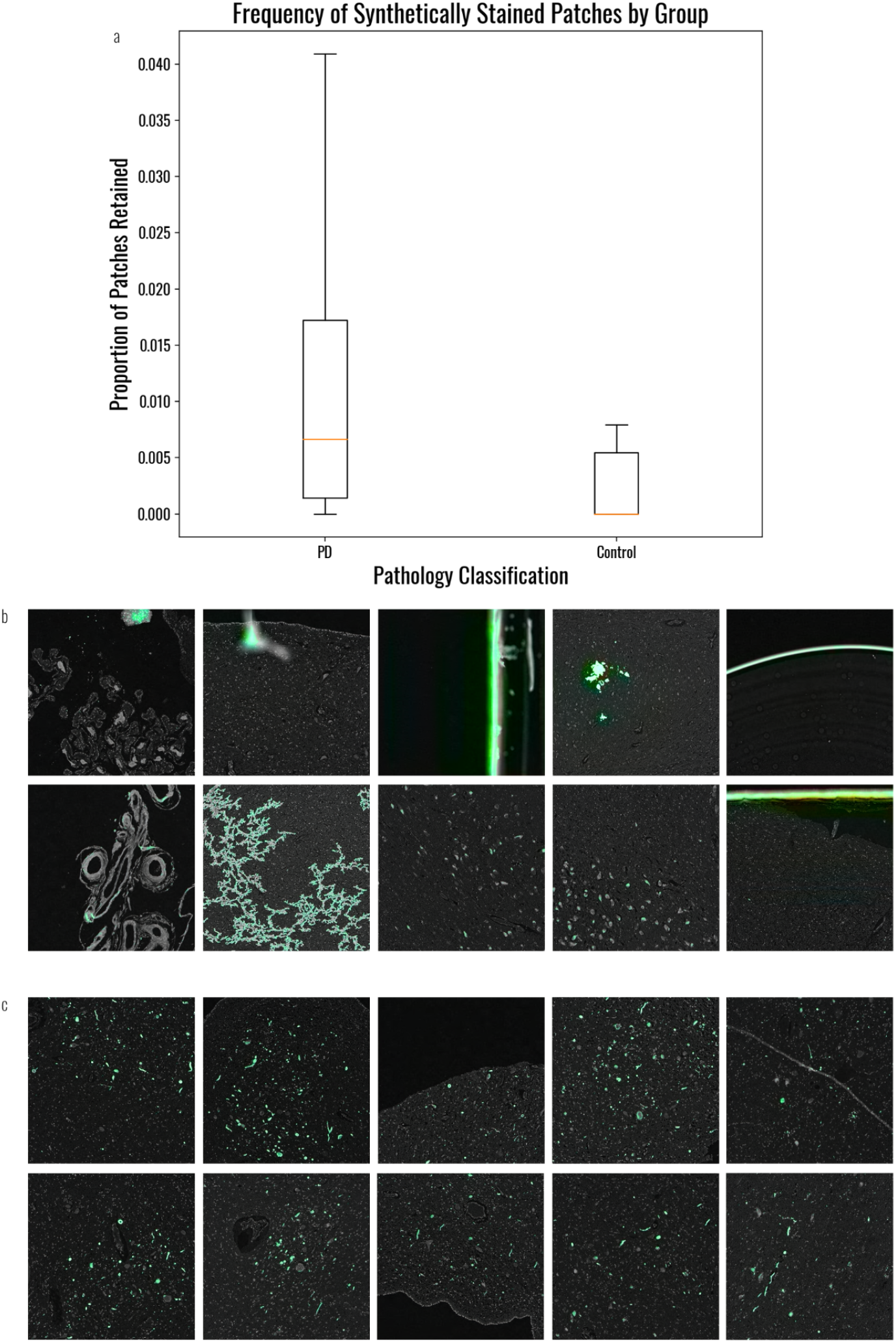
Frequencies of patches retained after filtering with examples of the visual quality of those patches. **a.** Patch densities for the different groups, demonstrating significantly more patches passing filtering in the PD group. Box plots show quartiles of the distribution of patches retained for each case for slide 17; outliers are discarded. Medians are marked in orange. **b.** Examples of the erroneous retained patches from both the control group and PD groups, these include: neuromelanin, edges of the slide, filaments and other foreign bodies, and darkly pigmented regions such as some cell nuclei; **c**. Examples of regions containing *α*-syn protein stained in patches from the PD group, including peripheral cell bodies.

### PDNet Training & Testing Results

Model training took approximately 10 hours to complete 500 epochs. Evaluation took less than 1 minute of elapsed time using the 25% validation slide cut of the slide dataset. Training, validation loss, and accuracy as a function of epoch can be seen below in Figure 3.5, along with the ROC for individual patch classification. Table 3.1 below shows signal-detection performance statistics (as used by Signaevsky et al. (2022)); we selected the best F1 threshold (confidence≅0.5±0.0001) to compare with expert raters from Signaevsky et al. (2022) for Braak 1 staging. All metrics are calculated as a mean of all batch statistics (error ±0.13%).

**Figure 3.5.**
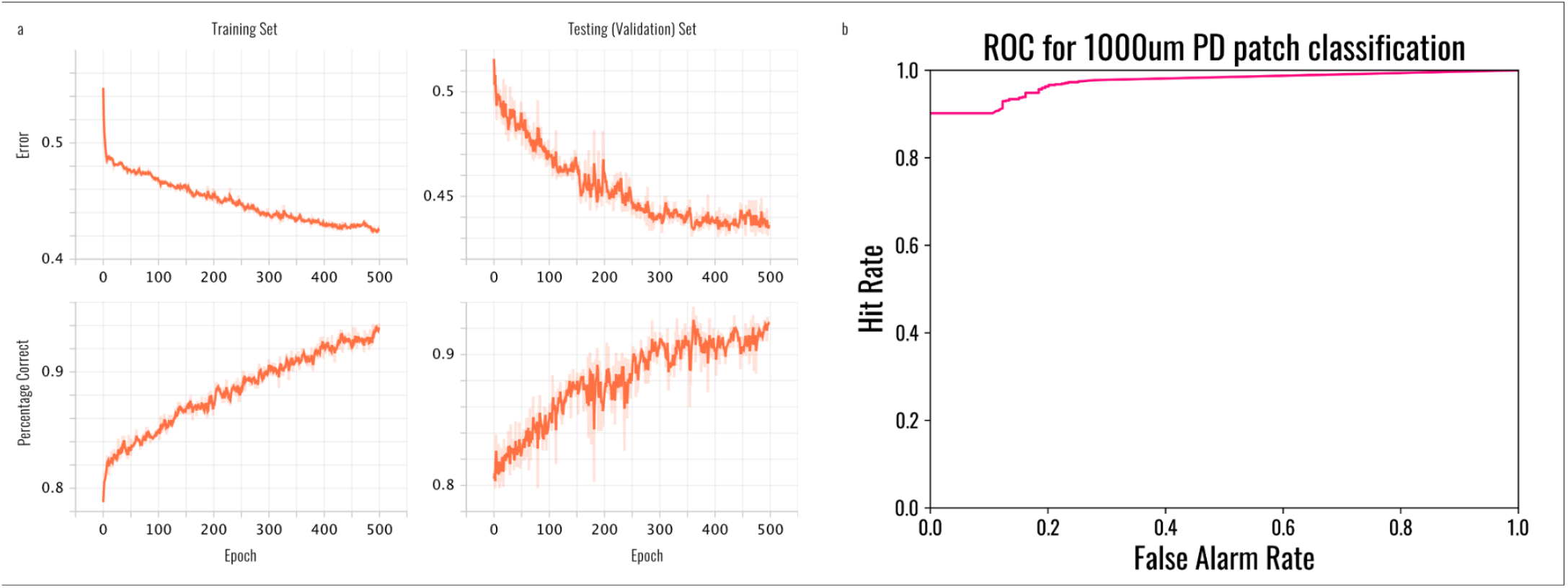
Performance characteristics as function of epoch and final binary classification as a function of model confidence. **a**. Training and validation loss (labelled Error) as a function of epoch; below, percentage correct for binary classification (confidence > 0); **b**. Receiver operating characteristic for binary classification of patches using a sliding threshold (confidence > T), where T is between 0 and the maximum confidence. Hit rate is capped for false alarm rates >.1, as no lower hit rate was observed than .9, due to the aggressive loss function used during training.

**Table 3.1.** PDNet performance characteristics per patch, with per case comparisons for clinicians, (both on Braak 1+ Staging).

| System | | Sensitivity<sup>3</sup><br>$tp / (tp + fn)$ | Specificity<sup>4</sup><br>$tn / (tn + fp)$ | Precision<sup>5</sup><br>$tp / (tp + fp)$ | F1<sup>6</sup><br>$2 * P * R / (P + R)$ | Accuracy<sup>7</sup><br>$(tp + tn) / N$ |
| --- | --- | --- | --- | --- | --- | --- |
| PDNet | | **0.93** $\pm 0.01$ | **0.88** $\pm 0.01$ | **0.88** $\pm 0.01$ | **0.91** $\pm 0.01$ | **0.90** $\pm 0.01$ |
| Expert Raters | Signaevsky et al. (2022) | 0.59 $\pm 0.16$ | **0.88** $\pm 0.30$ | **0.94** $\pm 0.13$ | 0.71 $\pm 0.12$ | 0.66 $\pm 0.13$ |

These performance statistics show higher accuracy, and sensitivity at detecting PD cases than expert raters. Expert raters are slightly more specific and precise, if one considers the error reported elsewhere as an overestimate.

Processing time (elapsed time on the development machine) for each phase of the processing pipeline can be seen in Table 3.2. Rows indicate successive stages.

**Table 3.2.**
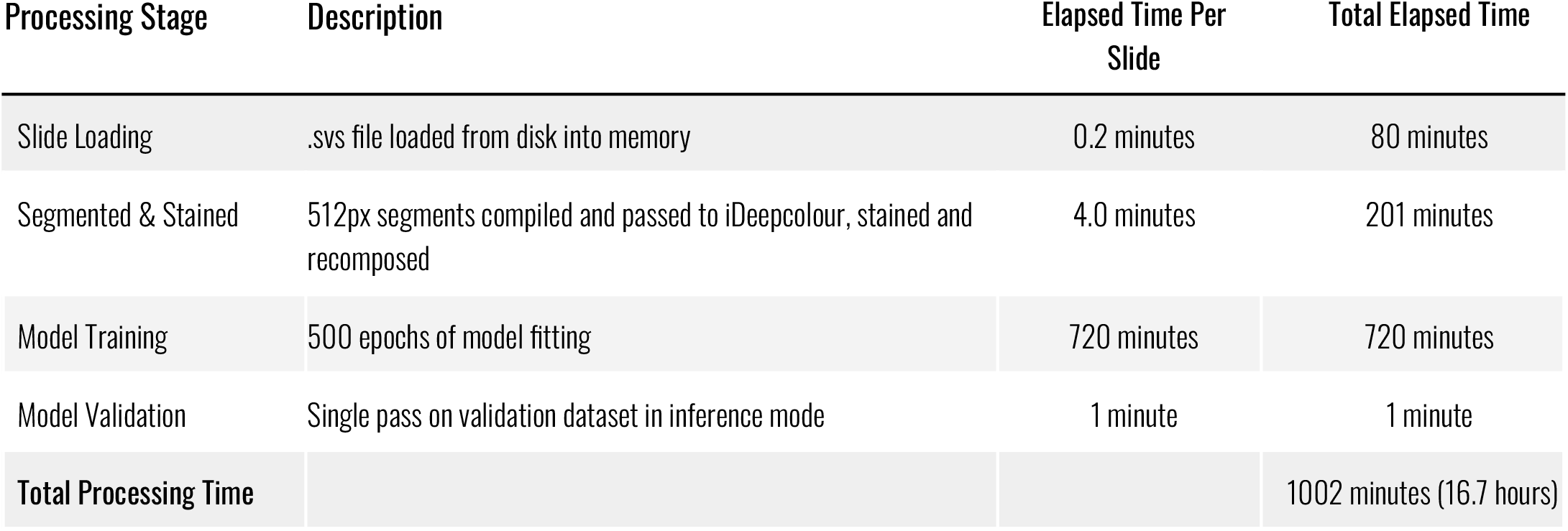
Processing time (elapsed time) on System (AMD Ryzen, Geforce RTX 3090) for each phase of processing for Slide 17 (dorsal motor nucleus of the Vagus). *See Materials for hardware description*.

## 4 Discussion

### Synthetic Staining & Post Processing

The synthetic staining process demonstrated the addition of useful information via two main mechanisms: filling in *α*-syn bodies in a texture-aware fashion, that would not be fully filled by a simple mask, and failing to fill highly textured bodies which were erroneously masked; this is evidenced by the smoothly filled outputs, and the jagged nature of the mask. The system (with the default chosen parameters) is sensitive to *α*-syn masking, evidenced with significantly more patches being stained in the PD group compared to controls (as shown in the box plots in Results-more patches remain after filtering). We identified that the median amount of staining per slide in the control group was None (the median line in the box plots, showing the midpoint in the control group distribution was 0 patches retained), with a median of 3 patches passing through in the PD group-proving the differential preference for *some* differentiating quality in the PD group slides (assumed here to be *α*-syn).

The thresholds used for segmentation and filtering were chosen as a reasonable starting point, however a parameter search should be performed to determine the optimal pruning parameters, as well as the optimal initial masking values for the synthetic staining process, and the optimal window sizes, as well as other hyperparameters. It may be the case that different search parameters show a greater difference between groups, and finding the threshold that collapses the difference would be informative; as it would describe fundamental qualitative and quantitative image differences between the sample groups.

That said, we have rejected the null hypothesis (that being there is no measurable difference between the synthetically stained outputs between the control and PD groups), and can confirm that we have an automatic staining algorithm for *α*-syn protein. This algorithm, while only benchmarked on the dorsal motor nucleus of the Vagus due to time considerations, does stain successfully on each staging region, with uniform threshold configuration parameters-as confirmed by visual inspection of stained slide examples of each site, by expert raters in the project group.

Both PD and control groups exhibit erroneously stained patches (such as the edge effects of the slide housing and foreign debris discussed previously). While a thorough investigation of these has not been conducted, it appears that the 0.5% −1.0% of patches that pass through in the control cases are not anatomically related. This suggests that with system configuration, such as the integration window size, the amount of staining accepted, and removal of the slide surrounds, cleaner separation between the PD and control group could be achieved without the need for more advanced statistical analysis.

Reported here is patch acceptance (passthrough) or rejection (filtered) frequency. Once a thorough evaluation of the staining parameters has been established, the density of *α*-syn of each slide should be directly measurable (currently a patch with very high staining contributes the same information as one that has just passed the filtering threshold). This will have research implications on both binary and categorical classification of the PD groups with respect to staging.

The extent of the usefulness of these synthetically stained outputs to visual neuropathological diagnosis of slide materials has not been examined here. Initial indications are that the *α*-syn regions are easily visible on desktop displays at a regular viewing distances (when the slide subtends ~12° of visual angle), while the raw slide images suffer from insufficient visual contrast between the staining chromaticity against the background. That said, the ability to search for a *known* chromatic signal (that is, a synthetically stained signal), makes ROI identification trivial. It is expected that as the aforementioned parameters are honed, this region identification will become more accurate and more useful.

### PDNet Results

We can conclude that there is sufficient separation between the filtered PD and control groups, and that those differences can be learned using supervised learning. This is shown by the ability of PDNet (the model) to train, and classify PD patches with reasonably high sensitivity and specificity in a binary detection task. This is demonstrated by ~93% of 1000µm PD patches being detected by the algorithm, with a monotonic increase in true positives and false alarms after ~90% true positives have been detected (see ROC in Figure 3.5 for illustration).

It is important to note that these specifications are not meant to be prescriptive for an objectively correct and optimal system design, but these results indicate that a design approaching the optimal system may be arrived at empirically. The absolute accuracy of the system, while high, should be taken in the context that it is the last processing step in a chain of under-optimised preprocessing and filtering stages. It would be reasonable to expect that should the optimal prefiltering parameters be found, that binary and categorical classification of PD patches should be considerably more sensitive and specific. Moreover, it is evident from the loss and accuracy plots that the model had not fully converged, and it is expected that with a hyperparameter search, a more accurate classifier could be trained more quickly. That being said, this model demonstrates that the assumptions regarding the dataset are correct, and that synthetically stained bodies within the PD group are different in quality to those of the control group, and those differences can be expressed in terms of convolutional filters. Moreover, those filters can be determined through gradient descent, using binary cross-entropy loss. This is also corroborating evidence that the preprocessing stages are effective in highlighting differences between the groups, or at least preserving them.

System confidence scores appear to be meaningful. This is evidenced by the fact that as the threshold for acceptance of the output confidence (C) increases the false positive rate decreases, and as the acceptance threshold decreases the true positive and false positive rate rise monotonically (see the ROC in Figure 3.5 the line never goes back down once it has increased). Therefore, in a correctly configured, final system, the confidence score should reflect a measure of pathology. As the confidence score increases, the certainty of both categorical and binary classification should increase and correlate with the pathologist/expert judgement.

Case level classification on the basis of slides of the dorsal motor nucleus of the Vagus (slide index 17 for each case in our dataset) is not discussed here and is the subject of further investigation. With the prevalence of patches within the PD group in conjunction with a conservative threshold, case level judgments should be viable, once the preprocessing stages have undergone refinement for erroneous patch staining. The current model is both significantly more accurate than the most comparable *in silico* model using biopsies (as in Sinaevsky et al., 2022), and the associated expert rater performance. A combination of the sensitivity of the model to find ROIs, and the expert precision (~6% higher than the model), this processing pipeline should offer an enhancement in the current diagnostic workflow. As described elsewhere (Attems et al., 2021), there is significant variability in the acceptance for particular PD staging, and therefore a rigorous analysis of case level staging and binary classification, and confidence scores should be evaluated.

A core contribution of this staining and classification process is the ability to automatically filter regions of interest for statistical analysis, which marks a major contribution to the neuropathological diagnosis of PD. We demonstrate that an automatic technique can load, process and identify regions of interest to the pathologist, and perform automatic binary classification, with high accuracy.

## 5 Next Steps

While there are innumerable possible next steps for utilising this technology we have included those of particular interest and focus for the project team, including those which we believe can be achieved within months of development effort and with maximum applicability to research.

If we consider no further development on the processing pipeline, the obvious next steps are to examine the parameters for each processing stage and to make those stages as widely applicable to other datasets and as sensitive as possible to *α*-syn signatures. This will allow high accuracy automatic ROI identification and classification beyond the current dataset and allow more feature rich signatures to pass further down the processing chain.

An examination of the viability of faster (to both implement and execute) and more widely understood techniques may lead to a widely accepted protocol, for automatic *α*-syn identification and PD classification. Such an examination of computationally intensive algorithms is desirable.

These techniques could also be applied to the identification of other target proteins, such as hyperphosphorylated tau, and amyloid beta. A particular focus should be on conjoined probability so that the nuanced pathological markers between dementia with associated Lewy Bodies and PD can be better understood.

These techniques should be systematically applied to each anatomical slide site, with the algorithm’s confidence used to produce categorical classification of PD stage. Ideally this additional information could feed into the training stage to allow differentiation of the filter development and the pathological progression of PD. This could also include the ability to utilise multiple sites, or the entire case scan set, to perform classification.

## 6 Conclusion

This project set out to explore whether digitised slides from PD cases could be loaded into machine learning frameworks, and if so, whether some differentiation could be made between PD cases and controls. Polygeist have delivered a PoC, command line library, which is capable of automatically loading, staining, segmenting and classifying regions associated with PD with state-of-the-art accuracy. These results are still early stages, with significant development work required to move this research towards a high Technology Readiness Level. The fundamental hypothesis that *α*-syn can be automatically stained, detected and classified has been upheld. PoC identification performance approaches, and in terms of visual search, is superior to that of the human visual system.

## Glossary of Specific Terms

| Term | Description |
| --- | --- |
| (Alpha) α-syn | Alpha-synuclein, a constituent of Lewy Bodies. |
| CNN | Convolutional Neural Network, an implementation of neural networks commonly used in computer vision tasks that takes inputs and repeatedly convolves them with arrays of values (weights) to produce outputs of a desired dimensionality. |
| JND | Just noticeable difference, the point at which a system starts to produce differential outputs randomly; for example, at this point in a binary system the outputs would alternate 50/50 between 0 and 1. |
| LBDs | Lewy Body Diseases, those diseases associated with the presence of Lewy Bodies. |
| Mask | An array of binary values corresponding to pixel coordinates in an image. Where the values are true, the pixel coordinates are retained. |
| Patches | A region of an image starting at a given coordinate and spanning a number of pixels width and height; for example, a 32x32 pixel image, which contains 1024 pixels, can be subdivided into 16 patches spanning 8 pixels in width and height, each patch containing 64 pixels. |
| PD | Parkinson's Disease. |
| PNG | Portable Network Graphic, an image container that provides lossless compressed rasterised imagery. |
| PoC | Proof of Concept, often related to a Technology Readiness Level of 4. |
| ROC | Receiver Operating Characteristic, a curve which describes the proportion of hits to false alarms generated by a system for a given threshold on the outputs. |
| ROI | A Region of Interest, a specified area within imagery which contains objects or other signals to be detected or analysed. |
| Slides | Digitised imagery of sections of brain tissue. |
| TRL | Technology Readiness Level, a measure of the operational maturity of a technology. |

## Acknowledgements

This project was managed and delivered by the Home Office’s Accelerated Capability Environment (ACE), with funding from the NHS Transformation Directorate AI Lab Skunkworks programme. We would like to thank the following people for their support and contributions to the project including technical assurance, project planning, roadmapping and testing: Amadeus Stevenson, Charlie Coleman, Matthew Cooper, Oludare Akinlolu, and Giuseppe Sollazzo from AI Lab Skunkworks; Tom McGeoch, Tom Park-Paul, Natalie Haarer, and Zain Ul-Haq from ACE; Romel Gravesande from Parkinson’s UK and others from the Parkinson’s UK Brain Bank at Imperial College London.

## Footnotes

1 Krippendorf’s *α* measures the average level of agreement between raters, who are using a unit scale (like the level of disease present in a case) to score an observation. A high score indicates a high level of agreement.

2 Type I errors occur when one accepts that an observed difference in a dependent variable has been produced by interacting with levels of the independent variable (or group), when instead those observations were in fact produced by extraneous random effects (the null hypothesis). A *p* < .001 indicates that we should expect this once per 1000 experiments, if our test assumptions are applicable.

## References

Alafuzoff, I., Ince, P. G., Arzberger, T., Al-Sarraj, S., Bell, J., Bodi, I., … & Kretzschmar, H. (2009). Staging/typing of Lewy body related *α*-synuclein pathology: a study of the BrainNet Europe Consortium. Acta neuropathologica, 117(6), 635–652.

Attems, J., Toledo, J. B., Walker, L., Gelpi, E., Gentleman, S., Halliday, G., … & McKeith, I. G. (2021). Neuropathological consensus criteria for the evaluation of Lewy pathology in post-mortem brains: a multi-centre study. Acta neuropathologica, 141(2), 159–172.

Bankhead, P., Loughrey, M.B., Fernández, J.A. et al. (2017). QuPath: Open source software for digital pathology image analysis. Sci Rep 7, 16878. https://doi.org/10.1038/s41598-017-17204-5

Beach, T. G., Serrano, G. E., Kremer, T., Canamero, M., Dziadek, S., Sade, H., … & Chahine, L. (2018). Immunohistochemical method and histopathology judging for the systemic synuclein sampling study (S4). Journal of Neuropathology & Experimental Neurology, 77(9), 793–802.

Bishop, C. M., & Nasrabadi, N. M. (2006). Pattern recognition and machine learning. (Vol. 4, No. 4, p. 738). New York: springer.

Braak, H., Del Tredici, K., Rüb, U., De Vos, R. A., Steur, E. N. J., & Braak, E. (2003). Staging of brain pathology related to sporadic Parkinson’s disease. Neurobiology of aging, 24(2), 197–211.

Dustler, M., Bakic, P., Petersson, H., Timberg, P., Tingberg, A., & Zackrisson, S. (2015, March). Application of the fractal Perlin noise algorithm for the generation of simulated breast tissue. In Medical Imaging 2015: Physics of Medical Imaging (Vol. 9412, pp. 844–852). SPIE.

Galasko, D. (2017). Lewy body disorders. Neurologic clinics, 35(2), 325–338.

Haykin, S. (2004). A comprehensive foundation. Neural networks, 2(2004, p. 173), 41.

He, Y., Hu, T., & Zeng, D. (2019). Scan-flood fill (SCAFF): An efficient automatic precise region filling algorithm for complicated regions. In Proceedings of the IEEE/CVF Conference on Computer Vision and Pattern Recognition Workshops (pp. 0–0).

Itti, L. & Koch C. (2001). Computational Modeling of Visual Attention. Nature Reviews Neuroscience 2(3):194–203

Law, A. M., Yin, J. X., Castillo, L., Young, A. I., Piggin, C., Rogers, S., … & Oakes, S. R. (2017). Andy’s Algorithms: new automated digital image analysis pipelines for FIJI. Scientific reports, 7(1), 1–11.

Lima, A. A., Mridha, M. F., Das, S. C., Kabir, M. M., Islam, M. R., & Watanobe, Y. (2022). A Comprehensive Survey on the Detection, Classification, and Challenges of Neurological Disorders. Biology, 11(3), 469.

McKeith, I. G., Galasko, D., Kosaka, K., Perry, E. K., Dickson, D. W., Hansen, L. A., … & Perry, R. H. (1996). Consensus guidelines for the clinical and pathologic diagnosis of dementia with Lewy bodies (DLB): report of the consortium on DLB international workshop. Neurology, 47(5), 1113–1124.

Melnikov, S., & Popov V. (2021). SlideIO Documentation. Retrieved from https://github.com/Booritas/slideio on 7 July 2022.

Mei, J., Desrosiers, C., & Frasnelli, J. (2021). Machine learning for the diagnosis of Parkinson’s disease: a review of literature. Frontiers in aging neuroscience, 13, 633752.

Munadi, K., Muchtar, K., Maulina, N., & Pradhan, B. (2020). Image enhancement for tuberculosis detection using deep learning. IEEE Access, 8, 217897–217907.

Olson, A. H. (2013). Calibration of Leica Scanscope AT2. Proceedings of the ICC Medical Imaging Working Group of the International Color Consortium. Retrieved from https://www.color.org/groups/medical/Olson.pdf on 7 July 2022.

Pointer, I. (2019). Programming pytorch for deep learning: Creating and deploying deep learning applications. (p. 58, 60). O’Reilly Media.

Redmon, J., Divvala, S., Girshick, R., & Farhadi, A. (2016). You only look once: Unified, real-time object detection. In Proceedings of the IEEE conference on computer vision and pattern recognition (pp. 779–788).

Signaevsky, M., Marami, B., Prastawa, M., Tabish, N., Iida, M. A., Zhang, X. F., … & Crary, J. F. (2022). Antemortem detection of Parkinson’s disease pathology in peripheral biopsies using artificial intelligence. Acta Neuropathologica Communications, 10(1), 1–14.

Szegedy, C., Ioffe, S., Vanhoucke, V., & Alemi, A. A. (2017, February). Inception-v4, inception-resnet and the impact of residual connections on learning. In Thirty-first AAAI conference on artificial intelligence.

Taha, A. A., & Hanbury, A. (2015). Metrics for evaluating 3D medical image segmentation: analysis, selection, and tool. BMC medical imaging, 15(1), 1–28.

Tan, M., & Le, Q. (2019, May). Efficientnet: Rethinking model scaling for convolutional neural networks. In International conference on machine learning (pp. 6105–6114). PMLR.

Yang, Y., Zhang, L., Du, M., Bo, J., Liu, H., Ren, L., … & Deen, M. J. (2021). A comparative analysis of eleven neural networks architectures for small datasets of lung images of COVID-19 patients toward improved clinical decisions. Computers in Biology and Medicine, 139, 104887.

Zhang, R., Zhu, J. Y., Isola, P., Geng, X., Lin, A. S., Yu, T., & Efros, A. A. (2017). Real-time user-guided image colorization with learned deep priors. arXiv preprint arXiv:1705.02999.

